# *segment_liftover*: a Python tool to convert segments between genome assemblies

**DOI:** 10.1101/274084

**Authors:** Bo Gao, Qingyao Huang, Michael Baudis

## Abstract

The process of assembling a species’ reference genome may be performed in a number of iterations, with subsequent genome assemblies differing in the coordinates of mapped elements. The conversion of genome coordinates between different assemblies is required for many integrative and comparative studies. While currently a number of bioinformatics tools are available to accomplish this task, most of them are tailored towards the conversion of single genome coordinates. When converting the boundary positions of segments spanning larger genome regions, segments may be mapped into smaller subsegments if the original segment’s continuity is disrupted in the target assembly. Such a conversion may lead to a relevant degree of data loss in some circumstances such as copy number variation (CNV) analysis, where the quantitative representation of a genomic region takes precedence over base-specific accuracy. *segment_liftover* aims at continuity-preserving remapping of genome segments between assemblies and provides features such as approximate locus conversion, automated batch processing and comprehensive logging to facilitate processing of datasets containing large numbers of structural genome variation data.

## Introduction

The first draft version of human genome was published in 2001[1]. In subsequent years, several new editions were released to perfect the quality of the genome assembly. The current version of human genome (GRCh38, UCSC hg38) was made available in 2013, with the latest revision (Grch38.p12) still containing more than 10 million unplaced bases [2]. Over the years, large numbers of genomic studies have been performed, generating data mapped to different versions of the reference genome. However, when performing genome analyses integrating data from multiple resources, it is imperative to convert all data to the same genomic coordinate system.

Two general methodologies are used for conversion between coordinates from different genome assemblies. The first approach is to re-align the original sequence data to the target assembly. This method could provide the best result but is very time consuming, and is not possible when the original sequence data is not available or does not consist of direct sequences (i.e. segmentation of array based data). Another approach is to convert the coordinates of genome data between assemblies by using a mapping file. This method, although bearing a side effect of minor information loss, for most applications provides a good balance between performance and accuracy.

Currently, three tools are in widespread use for the conversion between genome assemblies by coordinates: *liftOver* from University of California, Santa Cruz (referred as *UCSC liftOver* in the following article)[3]; *CrossMap* from Zhao[4]; and *Remap* from NCBI[5]. The *UCSC liftOver* tool exists in two flavours, both as web service and command line utility. It offers the most comprehensive selection of assemblies for different organisms with the capability to convert between many of them. *CrossMap* has the unique functionality to convert files in BAM/SAM or BigWig format. It generates almost identical results as *UCSC liftOver*, but is not optimised for converting genome coordinates between species. *Remap* provides for each organism a comprehensive list of major assemblies and the corresponding sub-versions. It can also perform cross species mapping, however, with only a limited number of organisms.

All those tools are efficient in coordinate conversion and provide almost identical results. However, as shown in figure 1a, challenges arise when dealing with genome segments that are not continuous anymore in the target assembly; there, these three tools take on different strategies. *CrossMap* and *UCSC liftOver* break the segment into smaller segments and map them to different locations. *Remap* keeps the integrity of the segment and maps the span to the target assembly.

**Figure 1.**
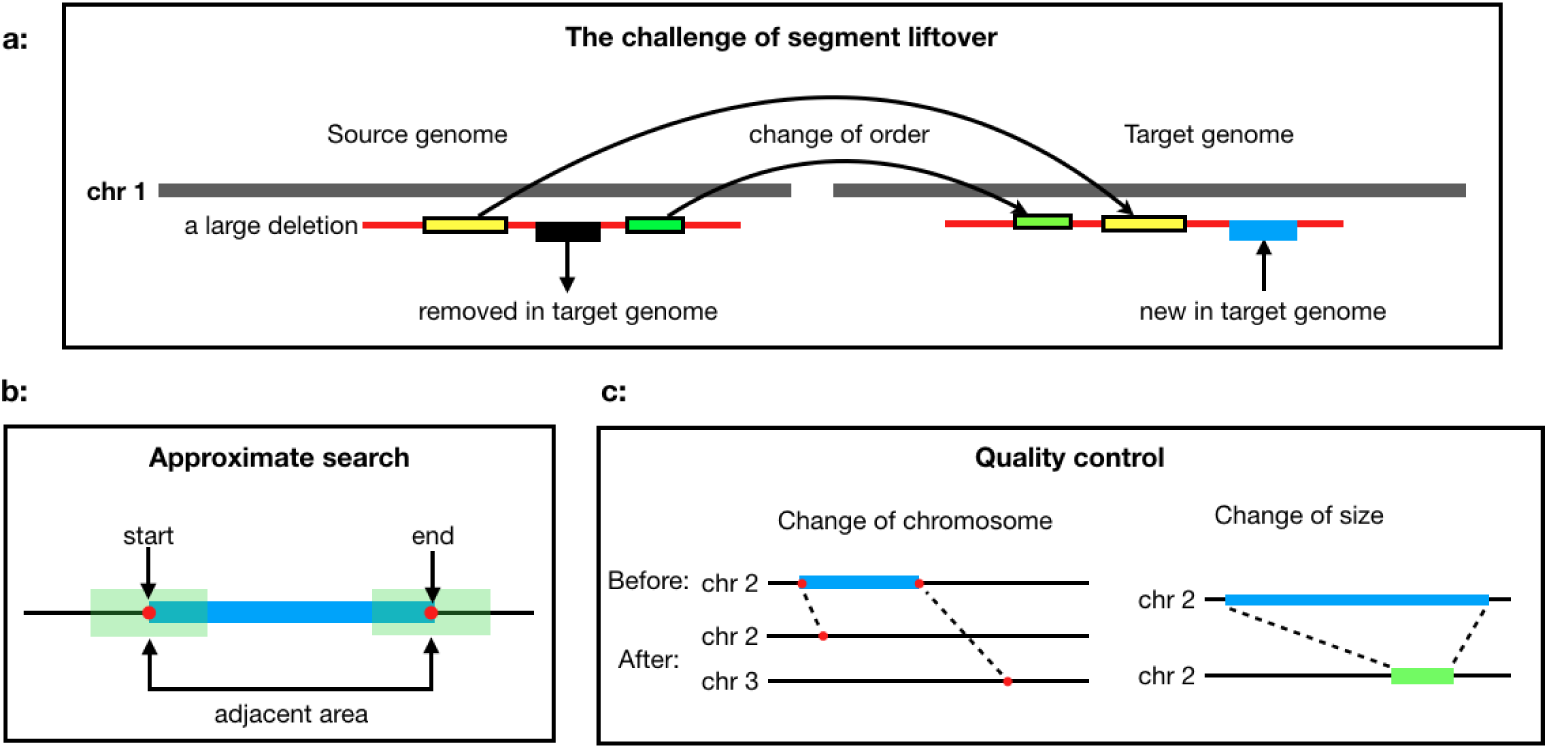
The challenge of segment liftover: (a) When lifting a segment to another assembly, the landscape of the segment may be affected by indels and copy number variations, but the overall span of the segment does not change significantly. (b) When the end positions cannot be converted by the *UCSC liftOver*, the nearby regions will be searched for convertible positions as approximation. (c) Quality control checks for changes of chromosome or size to make sure the segment is converted properly.

In research such analysis of copy number variation (CNV) data, where the quantitative representation of a genomic range takes precedence over base-specific representation, the integrity of a continuous segment indicates the proper conversion between assemblies, but may not be a direct outcome of current re-mapping approaches. Although *Remap* can convert contiguous segments, it only provides web service with submission limits, which is difficult to use for large scale or pipelined applications. The limitation to single input files is a general limitation of those tools, which precludes their direct use in comparative studies which may require to work with tensor hundreds of thousands of files, as indicated through our own projects and requests from the research community.

In this article,we introduce *segment_liftover*, a tool to perform an integrity-preserving conversion of genomic segments data between genome assemblies. It features two major functional additions over existing tools: First, re-conversion by locus approximation, in instances where a precise conversion of genomic positions fails; and second, the capability to handle any number of files and optional integration into data processing pipelines with supporting features such as automatic file traversal, interruption resumption and detailed logging.

## Methods

### Implementation

*segment_liftover* can convert both probe files and segment files at the same time or in separate runs. It starts from a structured directory or a list of files, then traverses and converts all files meeting the specified name pattern, and finally outputs to a designated directory. To convert a probe file, *segment_liftover* will first use the *UCSC liftOver* to convert the file, then apply an approximate conversion on probes that the *UCSC liftOver* failed to convert. To convert a segment file, *segment_liftover* will use the *UCSC liftOver* to convert the *start* and *end* positions of segments. A successful segment conversion needs to satisfy the following four criteria:

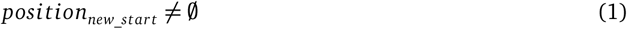

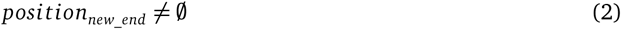

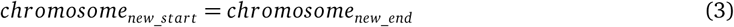

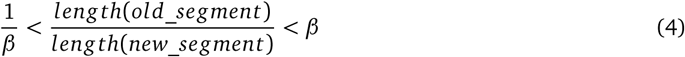

Where *β* controls the threshold of the length ratio and the default value is set to 2. If criteria (1) or (2) fails, *segment_liftover* will apply an approximate conversion; if the conversion still fails, it is reported as unconvertible. If criteria (3) or (4) fails, the conversion is reported as rejected (figure 1c). The reason of failure is recorded in log files.

When a position cannot be converted by the *UCSC liftOver, segment_liftover* will attempt an approximate conversion and try to find a convertible position in the adjacency (figure 1b). The range and the resolution of the search is defined by parameters *–range* and *–step_size*, respectively.

### Operation

The *segment_liftover* tool is implemented in Python. The package is available for both Linux and OSX. We recommend an installation using *pip* in a Python virtual environment. *segment_liftover* requires and depends on the *UCSC liftOver* program. A chain file, which provides alignments from source to target assembly, is also required. The chain files between common human assemblies (hg18, hg19 and hg38) are included in the program package. Chain files of other species and assemblies are available from the USCS Genome Browser. Figure 2 illustrates the work-flow of *segment_liftover*.

**Figure 2.**
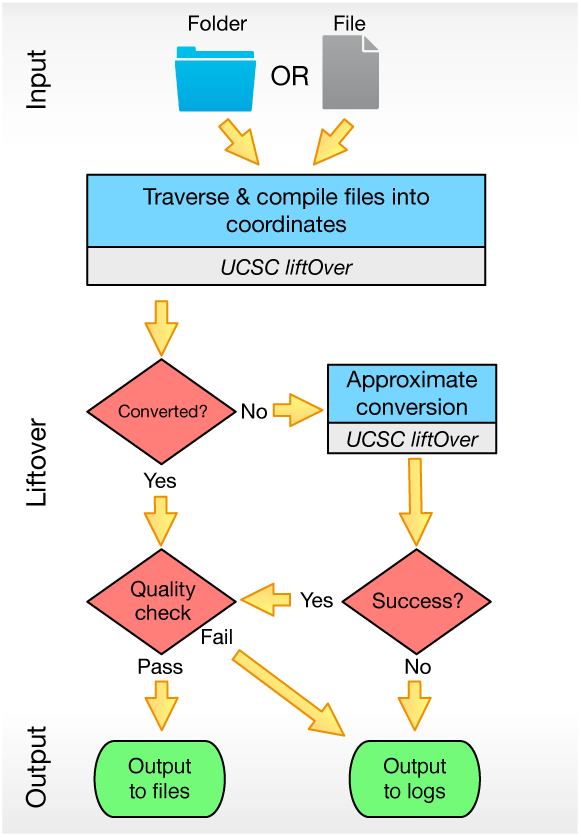
The workflow of *segment_liftover*: (1) It can take either a folder or a file containing the list of files as the input. (2) It will try to convert by approximation when *UCSC liftOver* fails to convert a coordinate. (3) The directory structure will be kept in the output folder and detailed log files are also available.

## Use Cases

### Converting arrayMap data from hg19 to hg38

In this section, we provide two examples of using *segment_liftover* to convert probes and segments, respectively. The two examples are part of the pipeline which updates the *arrayMap* database, a reference resource of somatic genome copy number variations in cancer[6], from human genome assembly hg19 to hg38. In the first example, we converted 44,632 probe files and 44,471 segment files from hg19 to hg38. The probe data were generated from nine Affymetrix genotyping array platforms, which currently only support annotations for hg19[7]. Circular binary segmentation (CBS) analysis (DNAcopy R-package) was used to infer copy number segments from log2 values of probes. The final segment files contain a list of genomic regions separated by their copy number values[8]. We ran the *segment_liftover* tool on a 12-core, 128GB RAM machine with 8 parallel processes. It took 42 hours to convert 44,632 probe files with 5.5 billion probe positions and 40 minutes to convert 44,471 segment files with 4.8 million segments.

Overall, more than 99.99% of probes and more than 99% of segments could be directly converted from hg19 to hg38 (Figure 3). As conversion of segments is more complicated and involves a quality control procedure to ensure the meaningfulness of the segment, it is expected to have a higher number of unconvertible segments than probes. As shown in Figure 4, the unconvertible regions are mainly around telomeres, centromeres, or other gene-sparse locations. In total, 38 genetic elements were found affected by the conversion (description in supplementary Table 1).

**Table 1.** Number of samples from nine platforms in use case examples

|  | all | 1000 samples |
| --- | --- | --- |
| CytoScanHD_Array | 2963 | 73 |
| CytoScan750K_Array | 173 | 2 |
| Mapping50K_Hind240 | 2699 | 58 |
| Mapping50K_Xba240 | 3303 | 72 |
| Mapping10K_Xba142 | 912 | 19 |
| Mapping250K_Nsp | 9738 | 223 |
| Mapping250K_Sty | 7561 | 184 |
| GenomeWideSNP_6 | 16570 | 359 |
| GenomeWideSNP_5 | 552 | 10 |

**Figure 3.**
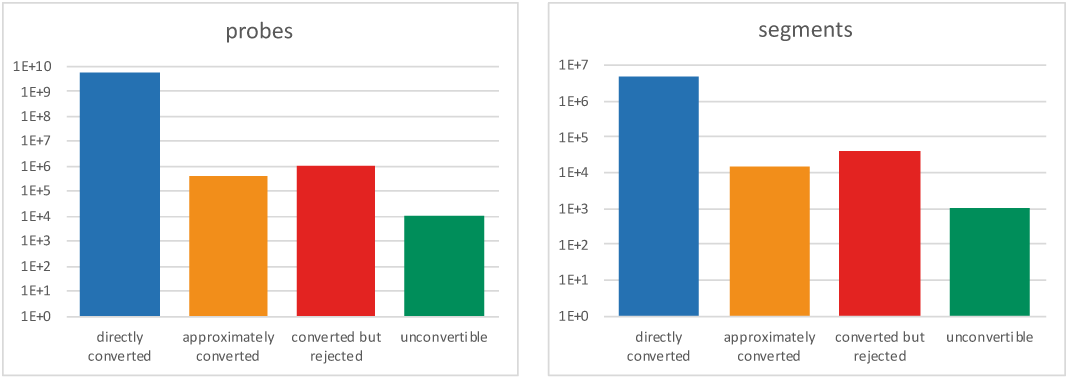
Compasion of conversion results in log 10. *Directly converted* is the sum of successful conversion from *UCSC liftOver*; *approximately converted* is the sum of successful reconversion in locus adjacency; *converted but rejected* is the sum of all rejections from quality control; *unconvertible* is the sum of everything that is not converted.

**Figure 4.**
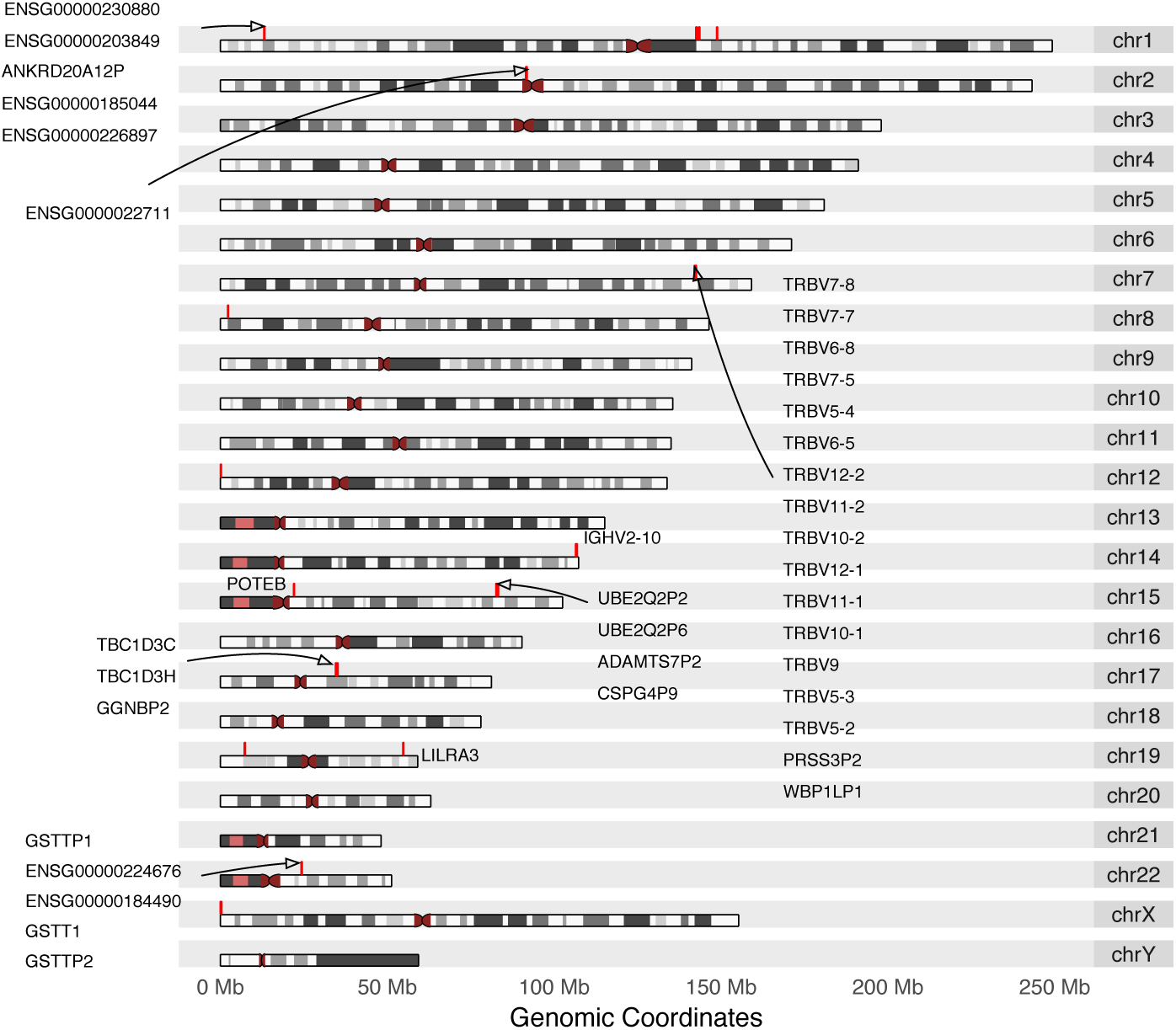
Genomic regions of unconvertible probe positions from human genome hg19. Unconvertible positions are marked red on the karyogram, annotated with HGNC symbol or ENSEMBL gene ID (if HGNC not available), retrieved from biomaRt_2.30.0.

### Comparison of different conversion strategies

In the second example, we compared the performance of different conversion strategies using 1,000 samples randomly drawn from the first example (Table 1). Copy number segments were generated using four different strategies(Figure 5): (1) segments in hg19 were generated from probes in hg19 using the aforementioned standard pipeline; (2) segments in hg38 were generated with probes converted from hg19 to hg38 using the standard pipeline; (3) segments converted from hg19 to hg38 with approximate conversion; (4) segments converted from hg19 to hg38 without approximate conversion.

**Figure 5.**
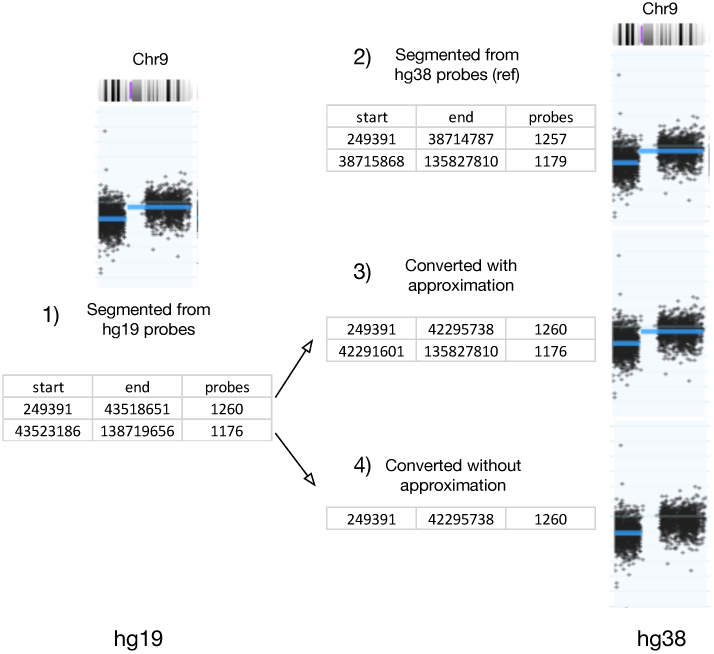
Chromosome 9 from GSM276858 with probe and segment data from hg19 coordinate(left) and hg38(right). Segments directly processed from hg38 probes are used as reference (top right). hg19 segments converted with approximation (middle right) and without approximation (bottom right) are used for comparison.

Table 2 shows the comparison of segments conversion in average between (3), (4) and (2) (complete table in supplementary Table 2). Exact segment matches are categorized as “perfect”; “minor difference” is defined the same as condition (3) of quality control; the rest of the segments are categorized as “significant difference”; “sum” is the total number of segments of a sample in average. By comparing the sum of *reference hg19, reference hg38* and *approximation*, it shows that the conversion result is very close to the result of the standard pipeline. The difference between converted and generated sums is much smaller than the difference between two generated sums of different genome versions. On average, the approximate conversion could rescue one additional segment per file. Finally, we zoom into a specific example on chromosome 9 in GSM276858 (illustrated in Figure 5). Because of the removal of probes from hg19 to hg38, the second segment will be lost without approximate conversion.

**Table 2.** Number of segments with or without approximation on average

|  | perfect | minor difference | significant difference | sum |
| --- | --- | --- | --- | --- |
| reference hg19 | na | na | na | 218.42 |
| reference hg38 | na | na | na | 215.04 |
| approximation | 198.18 | 6.24 | 10.25 | 214.67 |
| no approximation | 198.18 | 5.23 | 10.01 | 213.42 |

The two examples above demonstrated the efficiency and effectiveness of *segment_liftover* in processing large number of probe and segment files. It can provide conversion results that are similar to results generated from the standard pipeline. Moreover, with approximate conversion, the number of properly converted segments is slightly increased. In general, *segment_liftover* is able to provide reliable conversions and the ease of use.

## Summary

Translation between genome versions of sequencing data is a tedious but crucial task in bioinformatics. With the functionalities of automated batching, approximate conversion and segment conversion, *segment_liftover* can dramatically reduce the complexity and workload of such data processing. Furthermore, *segment_liftover*’s detailed logs of execution result provide an easy and clear foundation for follow up analysis.

## Software availability

pip version: https://pypi.python.org/pypi/segment-liftover

github: https://github.com/baudisgroup/segment-liftover

Software license: MIT

## Author contributions

BG developed the tool. QH worked on testing and data analysis. MB conceived the original concept. All authors contributed to the writing and revises of the manuscript.

## Competing interests

No competing interests were disclosed.

## Grant information

BG is recipient of a grant from the China Scholarship Council (CSC 201606620032). QH is supported through the University of Zurich’s “CanDoc” program.

## Acknowledgments

We thank Paula Carrio Cordo and the participants of the “Zurich Seminars in Bioinformatics” for helpful discussions.

